# A polygenic and phenotypic risk prediction for Polycystic Ovary Syndrome evaluated by Phenome-wide association studies

**DOI:** 10.1101/714113

**Authors:** Yoonjung Yoonie Joo, Ky’Era Actkins, Jennifer A. Pacheco, Anna O. Basile, Robert Carroll, David R. Crosslin, Felix Day, Joshua C. Denny, Digna R. Velez Edwards, Hakon Hakonarson, John B. Harley, Scott J Hebbring, Kevin Ho, Gail P. Jarvik, Michelle Jones, Tugce Karderi, Frank D. Mentch, Cindy Meun, Bahram Namjou, Sarah Pendergrass, Marylyn D. Ritchie, Ian B. Stanaway, Margrit Urbanek, Theresa L. Walunas, Maureen Smith, Rex L. Chisholm, International PCOS Consortium, Abel N. Kho, Lea Davis, M. Geoffrey Hayes

## Abstract

**Purpose:** As many as 75% of patients with Polycystic ovary syndrome (PCOS) are estimated to be unidentified in clinical practice. Utilizing polygenic risk prediction, we aim to identify the phenome-wide comorbidity patterns characteristic of PCOS to improve accurate diagnosis and preventive treatment.

**Methods and Findings:** Leveraging the electronic health records (EHRs) of 124,852 individuals, we developed a PCOS risk prediction algorithm by combining polygenic risk scores (PRS) with PCOS component phenotypes into a polygenic and phenotypic risk score (PPRS). We evaluated its predictive capability across different ancestries and perform a PRS-based phenome-wide association study (PheWAS) to assess the phenomic expression of the heightened risk of PCOS. The integrated polygenic prediction improved the average performance (pseudo-R^2^) for PCOS detection by 0.228 (61.5-fold), 0.224 (58.8-fold), 0.211 (57.0-fold) over the null model across European, African, and multi-ancestry participants respectively. The subsequent PRS-powered PheWAS identified a high level of shared biology between PCOS and a range of metabolic and endocrine outcomes, especially with obesity and diabetes: ‘morbid obesity’, ‘type 2 diabetes’, ‘hypercholesterolemia’, ‘disorders of lipid metabolism’, ‘hypertension’ and ‘sleep apnea’ reaching phenome-wide significance.

**Conclusions:** Our study has expanded the methodological utility of PRS in patient stratification and risk prediction, especially in a multifactorial condition like PCOS, across different genetic origins. By utilizing the individual genome-phenome data available from the EHR, our approach also demonstrates that polygenic prediction by PRS can provide valuable opportunities to discover the pleiotropic phenomic network associated with PCOS pathogenesis.

## Introduction

Polycystic ovary syndrome (PCOS) is the most common reproductive metabolic disorders, affecting 5-15% of reproductive age women worldwide [1]. The estimated cost of diagnosing and treating American women with PCOS is $5.46 billion annually as of 2017 [2, 3]. In addition to being a major cause of female infertility, the disease is a well-known risk factor for endocrine complications, such as type 2 diabetes, impaired glucose tolerance, and metabolic syndrome before age 40 [4]. Monozygotic twin studies of PCOS have suggested that PCOS is highly heritable (h^2^= ∼70%) [5] and the genetic architecture is polygenic with complex genetic inheritance pattern [6, 7]. Despite its clinical importance and high heritability, the underlying genetic etiology of PCOS remains incompletely understood. The phenotypic manifestations of PCOS are heterogeneous and exhibit considerable variation across race and ethnicity, further complicating the clinical diagnosis. Currently, it is estimated that up to 75% of women with PCOS remain undiagnosed in part due to varying diagnostic criteria from the National Institutes of Health (NIH), Rotterdam, or Androgen Excess Society, [8–12] which use different combinations of hyperandrogenism, ovulatory dysfunction, and/or polycystic ovarian morphology. Despite shared genetic risk across the criteria [13], the disagreement regarding PCOS phenotypic criteria presents a significant challenge for both clinical practice and research [14, 15]. The commonalities and differences between the phenotypic characteristics of PCOS may be better understood with an integrative observation of phenome-wide pleiotropies and co-morbidities.

Polygenic risk scores (PRS) built from well-powered genome-wide association studies (GWAS) have demonstrated operationalizing potential as biological risk predictors for patient stratification and risk prediction [16–19]. PRS represents the cumulative effect of common genetic variation summed per individual into a single risk score, providing an intuitive way to translate GWAS findings into clinically relevant information such as a patient’s risk of disease [20, 21]. From a precision medicine perspective, PRS hold significant promise especially for a multifactorial condition with complicated clinical manifestations, such as PCOS. However, several practical challenges remain in the equitable translation of PRS into clinical practice [22, 23]. For instance, most GWAS have been performed in samples of primarily European ancestry, resulting in PRS statistics that systematically perform worse in populations of different ancestry, including African ancestry populations. This underperformance is due to a combination of population-specific genetic effects that are undetected in a Euro-centric GWAS, and differences in the patterns of linkage disequilibrium (LD) between populations of differing biogeographic ancestry [24–27]. Thus, the evaluation of PRS from existing GWAS in both European and non-European ancestry samples is a critical step in setting priorities for equitable precision medicine initiatives.

The widespread deployment of Electronic Health Records (EHRs) and the availability of these multi-dimensional records enables evaluation of PRS in a research context that mimics a clinical hospital setting. Using these data, the predictive capability of PRS can be assessed regarding many possible diagnoses that can accumulate during an individual’s lifespan (i.e., the phenome). The eMERGE (electronic MEdical Records and GEnomics) Network is a nationwide consortium of multiple medical institutions that link DNA biobanks to EHRs [28], which is an important resource for determining the clinical utility of genomic findings, and enabling exploration of the range of phenotypes associated with genetic variation [29, 30].

The aim of this study is to systematically examine the utility of PRS derived from a GWAS meta-analysis by the International PCOS Consortium [13] for risk prediction across multiple ancestries and to further characterize the other EHR phenotypes that are clinically associated with PCOS genetic risk in both women and men. We first developed the integrative polygenic and phenotypic risk score (PPRS) for PCOS by combining the patient DNA genotype information and PCOS phenotypic elements from the EHR. Then we tested the predictive utility of the algorithm within European ancestry (EA) samples and further evaluated its performance in African ancestry (AA) and combined multi-ancestry (MA) participants which included EA, AA, and other ancestries. In addition, we performed a Phenome-Wide Association Study (PheWAS) of the PPRS for PCOS to identify the range of phenotypic indicators associated with PCOS and evaluated the predictive characteristics of PPRS to identify underlying PCOS pathophysiological pathways.

## Materials and Methods

### PCOS Polygenic Risk Score (PRS) Development

We obtained the full summary statistics of the largest meta-GWAS of PCOS through the International PCOS consortium and developed a PRS for PCOS [13]. **(Supplementary table 1)** The GWAS was conducted in 5,209 cases and 32,055 controls of EA women who were diagnosed according to either NIH or Rotterdam criteria. All variant positions were converted to GrCh37 and we excluded any entries with missing ORs or risk allele frequency (RAF) information. The RAF of each variant was calculated using PLINK [31], and we excluded the entries which RAF deviates more than 15% than our eMERGE data in order to ensure additional quality control (QC). PRSice software [32] was used to filter any correlated SNVs in pairwise Linkage Disequilibrium (LD) (r^2^ > 0.2) and constructed PRS for PCOS by summing the best-guess imputed genotype data of PCOS risk variants in each individual weighted by the reported effect sizes. We used eight different subsets of PCOS susceptibility SNVs to build the model based on p-value cutoff and compared for their predictive accuracy in the following validation step: 5×10^-8^, 5×10^-7^, 5×10^-6^, 5×10^-5^, 5×10^-4^, 5×10^-3^, 5×10^-2^, and 1 (All SNVs).

### PRS/PPRS Evaluation & PheWAS Discovery Cohort

Our cohort included genotypes and clinical diagnosis records of 99,185 individuals collected from 12 EHR-linked biobanks nationwide through the eMERGE consortium [29]. After identity-by-descent (IBD) analysis, we removed 8,019 related individuals that were not in canonical IBD position or genetically identical individuals near the origins (Z0 > 0.83 and Z1 < 0.1). The cohort was composed of multiple self-reported and 3^rd^ party observed ancestries and we defined them into three main genetic ancestral groups using the intersection of self-reported ancestries and principal component analysis (PCA) based k-mean clusters: European (71.7%), African (15.0%), and Asian (1.0%). We excluded any self-reported or genetically Hispanic participants for ancestry-stratified analysis for better homogeneity. Throughout this study, the first four principal components (PCs) were used to correct population structure, explaining over 17% of the variances among different genetic origins.

The phenome data of the participants were collected from the EHR including diagnostic records and basic demographic information. The data collection was performed under local institutional review board approval with informed consent from the patients. Diagnostic information was structured in the format of the International Classification of Diseases, Clinical Modification (ICD-CM) codes, in both 9th and 10th edition, and aggregated into a higher level of 1,711 phecodes for a standardized categorical analysis of diseases (Phecode map version 1.2) [33, 34]. We excluded 23 individuals under the age of 14, the clinically plausible age for PCOS diagnosis, which is defined as two years after the first menstruation. A demographic information of the 91,144 participants after filtering criteria is presented in Table 1.

**Table 1:**
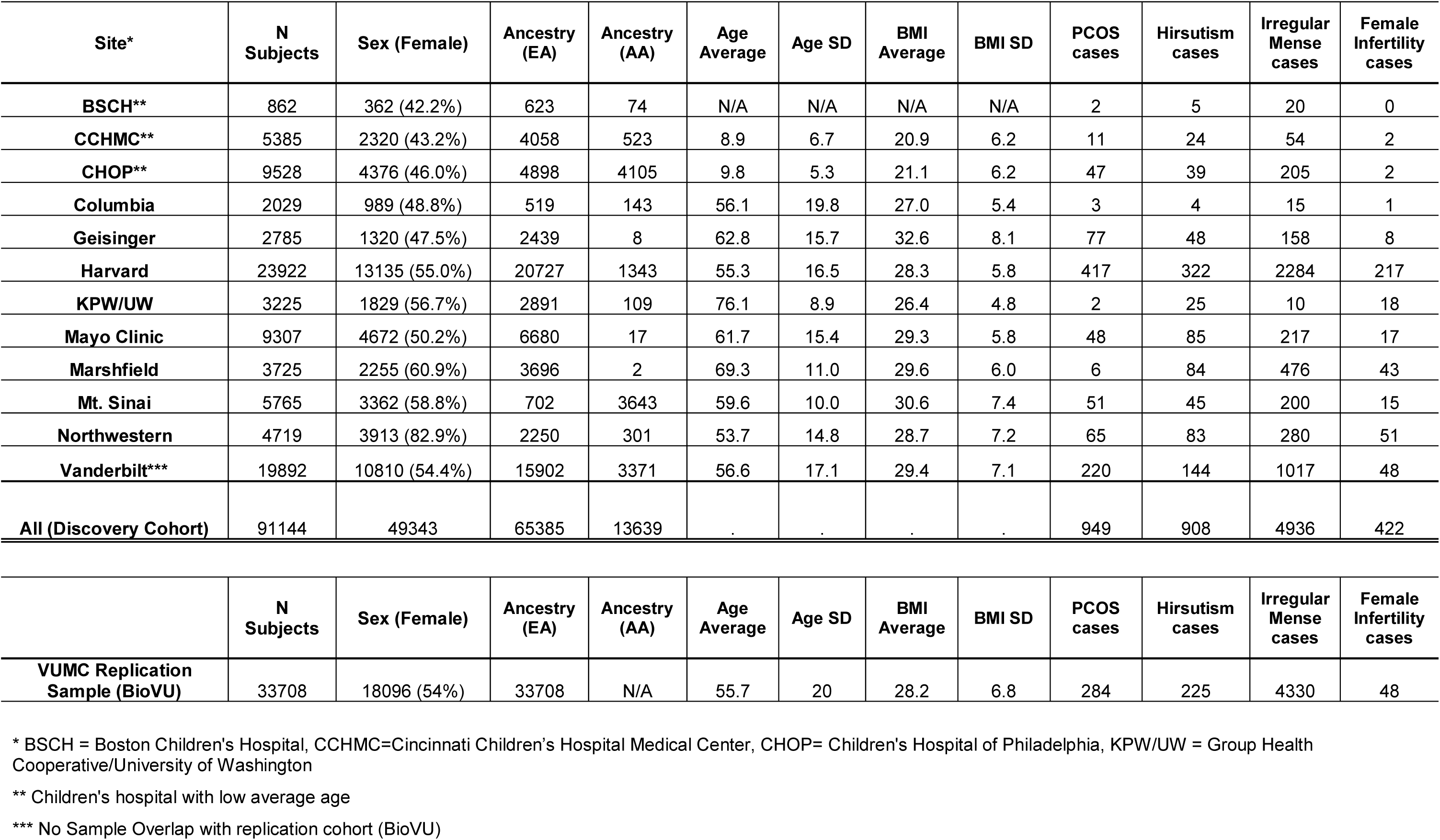
Demographic and clinical characteristics of discovery cohorts (eMERGE) and replication cohort (BioVU).

### Genotype data and Quality Control

The participants provided their saliva samples for genotyping, which were genotyped on 78 genotype Illumina or Affymetrix array batches from 12 medical sites.

We used the Michigan Imputation Server(MIS) [35] with the minimac3 missing genotype variant imputation algorithm to impute missing genotypes in our sample based on the Haplotype Reference Consortium (HRC1.1) which includes ∼65,000 individuals of diverse ancestry [36]. The imputation resulted in a genome-wide set of ∼40 million SNVs. We filtered the poorly imputed genetic variants with the r-squared imputation quality threshold (mean variant r-square) less than 0.3, minor allele frequency (MAF) less than 0.05 and genotype call rate lower than 95%, which resulted in 5,760,270 autosomal polymorphic variants for subsequent analysis. The detailed data collection and QC report for the eMERGE network is reported elsewhere [29].

### Validation of Polygenic Risk Score

#### A. Predictive ability of each prediction model with different PRS

We performed logistic regression analysis to demonstrate the prediction ability of PRS for PCOS diagnosis in the female population of three different genetic racial cohorts: European (n=33.869), African (n=8,198), and the entire admixed cohort (n=49,365). Each cohort was randomly divided into 75% training and 25% testing set to separately calculate the regression statistics and out-of-sample prediction error. Using generalized linear model, the residuals of PRS after covariate adjustments (first four PCs, sites) were obtained and scaled to build the logistic regression model in the training set. Regression coefficients and p-value of PRS variable, and pseudo-R^2^ of the eight different PRS models were measured.

We applied the regression model built out of the training set to measure out-of-sample performance in the testing dataset. We predicted the individuals as ‘PCOS cases’ if their fitted scores are higher than the average fitted score and calculated the accuracy by comparing with their actual diagnosis records of PCOS. The overall accuracy, sensitivity, specificity of each model were measured and structured through confusion matrix. The area under the receiving operating characteristic (ROC) curve, AUC, was also measured for classifier performance of each model.

#### B. Stratification ability of each prediction model with different PRS

To evaluate the phenotypic stratification ability of PRS, we divided the cohort into ten quantiles based on PRS of each individual and compared the average phenotypic values (e.g. proportion of PCOS diagnosed patients, body mass index (BMI), PRS) among the groups. The proportion of PCOS patients in each quantile, average PRS values, and average BMI measures of each individual were analyzed. We also performed independent t-test to assess if the average PRS score differences between PCOS cases and controls were statistically significant.

#### C. Performance improvement by the PRS variable

To estimate the performance of the PRS variable, we built a null regression model without the PRS variable for PCOS prediction (PRS model vs. Null model).The incremental pseudo-R^2^ by McFadden’s [37] were calculated between the PRS models and the null logistic regression only with first 4 PCs and site variables. The analysis of variance (ANOVA) was performed to examine how significant PRS variable impacts the PCOS diagnosis prediction model compared to the null model.

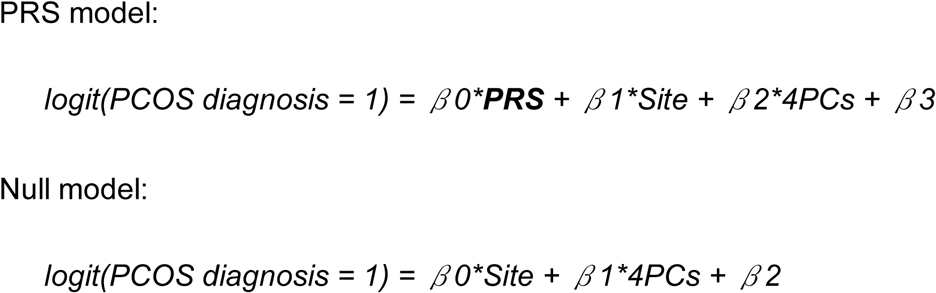

### Development of prediction algorithms with PRS and PCOS component phenotypes (PPRS)

We built an integrative polygenic and phenotypic risk score (PPRS) with PRS and PCOS component phenotypes in the EHR to maximize the utility of PRS for risk prediction. Additional dichotomous phenotypic variables to each individual from their EHR diagnosis records: hirsutism (ICD9 code 704.1, ICD10 code L68.0), irregular menstruation (ICD9 code 626.4, ICD10 code N92.6), and female infertility (ICD9 code 627, ICD10 code N97.0) were selected, all of which are well-established clinical components of PCOS. A total 908 individuals with hirsutism, 4,936 individuals with irregular menstruation, and 422 individuals with female infertility ICD diagnosis codes were identified in the eMERGE consortium database.

Firstly, the logistic regression adjusted for first four PCs and sites were examined for their effect coefficients and variable p-values. Psuedo-R^2^ of each model was calculated for measuring the improvement over the normal PRS model. ANOVA between the integrative model and normal PRS model were examined.

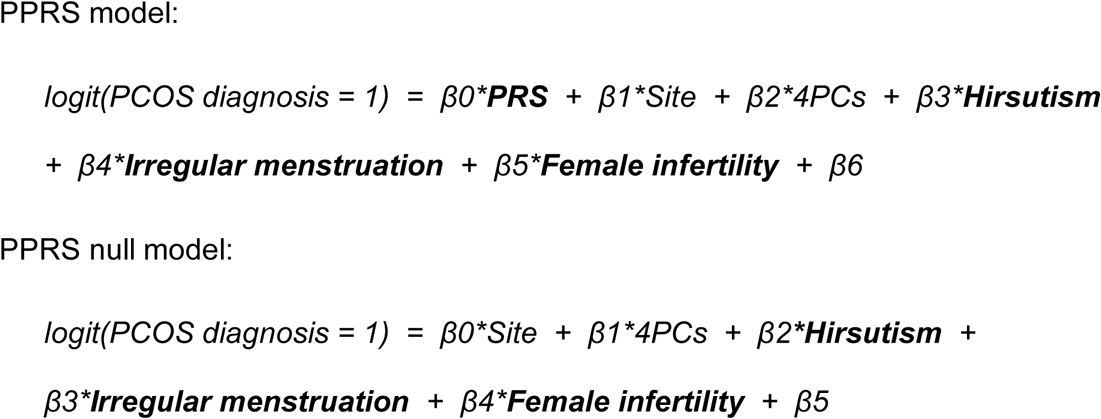

### Phenome-wide analysis

To investigate the potential pleiotropy of PCOS, PCOS components, and other diseases in the EMR phenome, we selected the best performing PRS model that presented a validated predictive accuracy and stratification ability across ancestries based on the examination results above. PheWAS was performed on the mapped 1,711 representative EHR phenotypes with a minimum of 30 case patients from the discovery cohort of 91,144 participants after QC criteria. Case group for a given phecode is defined by the presence of at least one assignment of the corresponding ICD codes from EHR as defined in the phecode map v1.2. Controls for each phecode are defined by the absence of the same ICD codes that defined cases and the absence of clinically related phenotypes. Based on the assumption that a participant with higher PCOS-PRS conveys greater genetic risk, our main sex-stratified PheWAS interrogated the comorbid networks of high-risk predictive phenotypes for PCOS (**PheWAS-1**). 49,343 female participants and 41,669 male participants were used for the analysis. Logistic regression was used adjusting for genotyping site and the first four PCs of ancestry to correct for population stratification in the MA cohort [*logit (Clinical Phenotype = 1 | PRS, Site, 4PCs) = β0 + β1*PRS + β2*Site + β3*4PCs*].

In this study, phenome-wide significance refers to either (1) the Bonferroni corrected threshold of p-value=2.9×10^-5^ adjusting for multiple testing, which is determined by using the p-value of 0.05 divided by the 1,711 phenotypes interrogated, or (2) the false discovery rate (FDR) significance of 0.05, which is a popular alternative threshold to the stringent Bonferroni correction in reporting PheWAS. Manhattan PheWAS plots of - log10(p-value) were generated for visual inspection of significant clinical traits. All the analyses were performed in the R statistical software environment (ver 2.1.2).

### Sensitivity Analysis

We performed several comparative PheWAS in an effort to interrogate different phenome-wide aspects of the PRS in clinical phenome.

Firstly, to distinguish secondary or symptomatic phenotypes derived from the PCOS-diagnosed patients, we removed the clinical diagnosis records of the 949 individuals with PCOS (phecode 256.4, ICD9 256.4 and ICD10 E28.2) and performed the same PheWAS analysis. **(PheWAS-2).** Additionally, to gauge the contrasting performance of polygenic prediction over a single-variant approach, we performed traditional PheWAS of each genome-wide significant susceptibility loci (p-value < 5×10^-8^) for PCOS (RAF > 0.05). This analysis aims to compare the clinical phenotypes associated with the cumulative effects of multiple genetic variants on PRS versus a single genetic signal generated by an individual PCOS susceptibility locus. Among 113 genome-wide significant loci (p-value < 5×10^-8^) for PCOS, (**Supplementary Table 1**) we filtered the entries with MAF > 0.05 and genotype call rate > 0.90 in our discovery cohort and MAF > 0.01 in summary statistics. 85 SNVs were selected and used for the subsequent PheWAS analysis (**PheWAS-3**).

### PRS PheWAS Replication

To confirm the predictive performance of our PRS algorithm and its effect on clinical phenome, replication analyses were performed at Vanderbilt University Medical Center on an independent genotyped sample of 33,708 European descent individuals (BioVU). The participants were genotyped on the Illumina MEGAEX platform (∼2 million markers) and we applied filters for individual call rates < 98%, batch effects (p-value < 5 x 10^-5^), heterozygosity (|Fhet| > 0.2), and sample relatedness (pihat > 0.2). After imputation with 1000G reference panel, we excluded any genetic variants with missingness > 0.02, certainty < 0.9, or imputation info score < 0.95. The genetic ancestry of the samples were restricted to only EA, based on comparison to 1000G European population and a K-means clustering definition. The final samples included 33,708 individuals of European descent genotyped on 5,550,390 SNVs. Using the same PRS generation methodology in discovery samples, we took the identical phenome-wide approach to identify the associated phenotypic networks with PRS among the replication samples. Logistic regression was used adjusting for first four ancestry PCs.

## Results

### Polygenic risk scores for PCOS are normally distributed in European and multi-ancestry participants

A total of 5,760,270 autosomal single nucleotide variants (SNVs) were considered for the PCOS-PRS construction, which displays the genetic architecture of effect size (beta) by risk allele frequency (RAF) presented in Figure 1. There was a significant negative correlation between RAF and effect size, which is generally anticipated in common quantitative traits and supports the use of methodology of PRS to explore the extreme of the common polygenic liability spectrum. According to the central limit theorem, PRS in a large population will show normality when the genetic architecture of the target trait is polygenic, i.e. produced by the addition of many genetic variants of small effect [38, 39].

**Fig 1.**
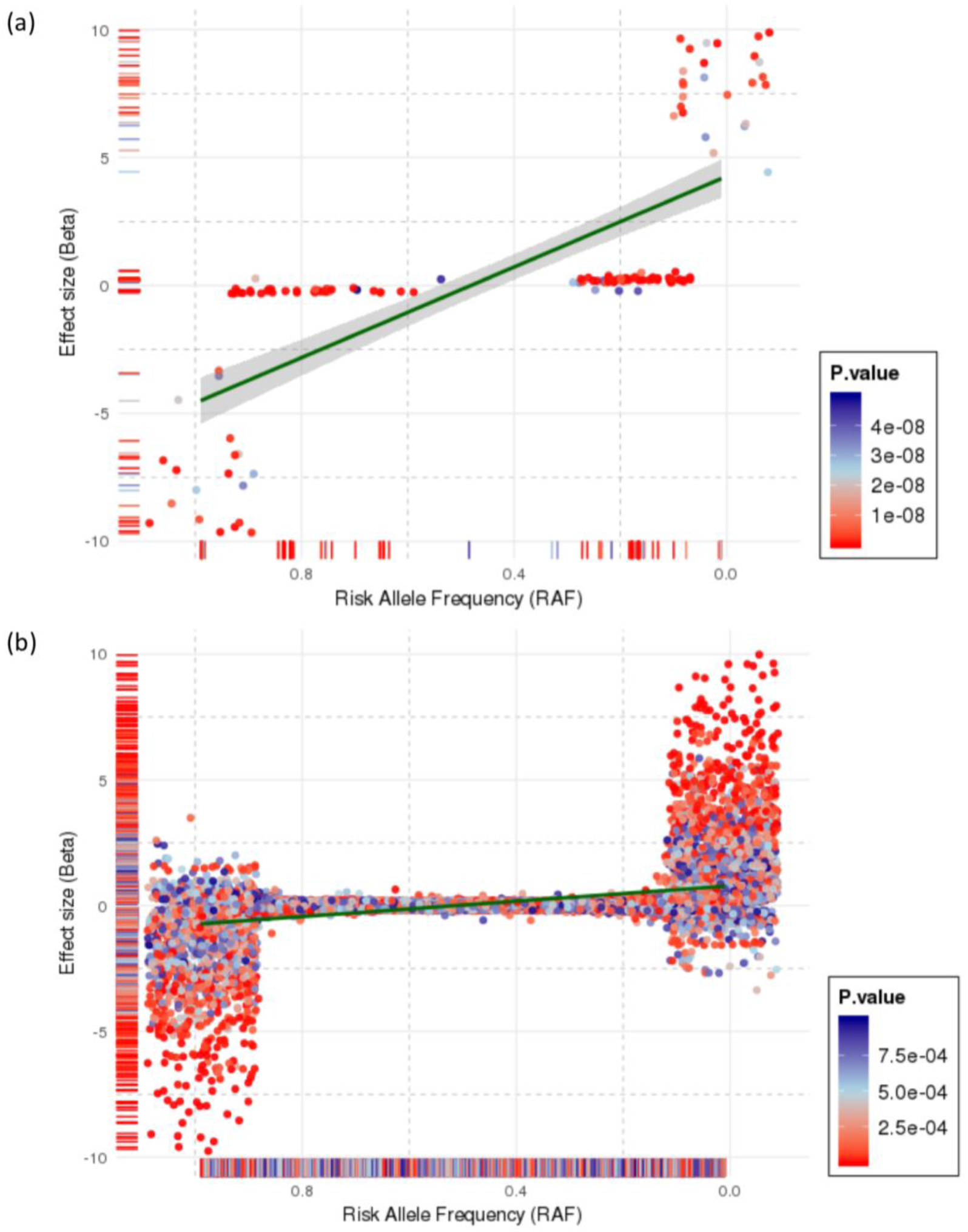
Effect distribution of PCOS susceptibility variants in samples from the International PCOS consortium by risk allele frequency. **(a)** The 120,340 PCOS autosomal SNVs with p-value < 0.05, and **(b)** the 139 PCOS genome-wide significant SNVs (p-value < 5×10^-8^). The dark green line and grey band around it are the linear regression fit and its 95% confidence interval, respectively, between risk allele frequency and effect size (beta).

PRS were calculated at eight different p-value cutoffs from the PCOS GWAS summary statistics (5×10^-8^, 5×10^-7^, 5×10^-6^, 5×10^-5^, 5×10^-4^, 5×10^-3^, 5×10^-2^, 1) for all the discovery eMERGE participants (n=91,144). Each set of scores were adjusted for participant site and first four PCs. All the polygenic scores were evaluated for their predictive performance in the female populations of EA (n=33,869), AA (n=8,198) and MA cohorts (n=49,365). The covariate-adjusted PCOS-PRS generally presented a normal distribution across each ancestry cohort (**Supplementary Figure 1**). PRS models with trimodal or skewed distributions (PRS p-value cutoff: 5×10^-7^, 5×10^-6^, 5×10^-5^), which may be a function of poor representation of risk variants across populations, were not considered for the subsequent phenome-wide analysis.

### Validation of PCOS PPRS in European ancestry participants

#### A. Predictive ability of each prediction model with different PRS

In the PRS prediction models using the training set of the female EA cohort (n=33,869 with 632 PCOS cases), all the coefficient p-values of the PRS variables are statistically significant except for two PRS models of SNVs with p-value < 5×10^-7^ and p-value < 5×10^-6^ that do not show PRS normality (**Supplementary Figure 1**). The average odds ratios (OR) of the significant PRS variable across EA was 1.13 (average SE=0.046) and the average pseudo-R^2^ value was 0.044, which indicates 4.4% of the phenotypic variances in the training sample could be explained by PRS (Table 2).

**Table 2:** Regression results of the PRS and PPRS models in PCOS prediction.

|  | PRS* |  |  |  | PPRS* |  |  |  |
| --- | --- | --- | --- | --- | --- | --- | --- | --- |
| PRS/PPRS<br>p-value<br>Cutoff | OR | Std.<br>Error | P | R<sup>2</sup> | OR | Std.<br>Error | P | R<sup>2</sup> |
| <u>European Ancestry</u> |  |  |  |  |  |  |  |  |
| 5E-08 | 1.14 | 0.047 | 4.76E-03 | 0.045 | 1.14 | 0.052 | 1.40E-02 | 0.232 |
| 5E-07 | 1.04 | 0.042 | 3.78E-01 | 0.043 | 1.04 | 0.046 | 3.89E-01 | 0.230 |
| 5E-06 | 1.08 | 0.039 | 6.41E-02 | 0.044 | 1.08 | 0.044 | 7.26E-02 | 0.231 |
| 5E-05 | 1.10 | 0.041 | 2.13E-02 | 0.044 | 1.10 | 0.046 | 3.59E-02 | 0.231 |
| 5E-04 | 1.13 | 0.045 | 6.12E-03 | 0.044 | 1.11 | 0.049 | 2.85E-02 | 0.231 |
| 5E-03 | 1.11 | 0.047 | 2.70E-02 | 0.044 | 1.08 | 0.051 | 1.45E-01 | 0.231 |
| 5E-02 | 1.16 | 0.048 | 2.11E-03 | 0.045 | 1.12 | 0.052 | 3.21E-02 | 0.231 |
| 1 | 1.15 | 0.049 | 3.13E-03 | 0.045 | 1.11 | 0.052 | 4.04E-02 | 0.231 |
| <u>Multiancestry</u> |  |  |  |  |  |  |  |  |
| 5E-08 | 1.16 | 0.038 | 1.15E-04 | 0.040 | 1.15 | 0.042 | 1.19E-03 | 0.228 |
| 5E-07 | 1.08 | 0.037 | 4.28E-02 | 0.038 | 1.09 | 0.038 | 2.99E-02 | 0.227 |
| 5E-06 | 1.09 | 0.037 | 1.60E-02 | 0.038 | 1.10 | 0.038 | 1.19E-02 | 0.227 |
| 5E-05 | 1.12 | 0.037 | 2.35E-03 | 0.039 | 1.12 | 0.038 | 3.67E-03 | 0.228 |
| 5E-04 | 1.12 | 0.038 | 1.88E-03 | 0.039 | 1.11 | 0.039 | 8.59E-03 | 0.228 |
| 5E-03 | 1.16 | 0.039 | 1.25E-04 | 0.040 | 1.13 | 0.041 | 2.54E-03 | 0.228 |
| 5E-02 | 1.20 | 0.039 | 5.03E-06 | 0.041 | 1.16 | 0.042 | 3.81E-04 | 0.228 |
| 1 | 1.22 | 0.040 | 5.33E-07 | 0.041 | 1.19 | 0.043 | 5.91E-05 | 0.229 |
| <u>African Ancestry</u> |  |  |  |  |  |  |  |  |
| 5E-08 | 1.14 | 0.090 | 1.42E-01 | 0.031 | 1.15 | 0.099 | 1.62E-01 | 0.211 |
| 5E-07 | 1.24 | 0.086 | 1.22E-02 | 0.034 | 1.30 | 0.093 | 4.63E-03 | 0.215 |
| 5E-06 | 1.25 | 0.086 | 9.80E-03 | 0.034 | 1.30 | 0.092 | 3.95E-03 | 0.216 |
| 5E-05 | 1.23 | 0.086 | 1.71E-02 | 0.034 | 1.27 | 0.093 | 1.08E-02 | 0.214 |
| 5E-04 | 1.19 | 0.088 | 4.38E-02 | 0.032 | 1.17 | 0.094 | 9.82E-02 | 0.211 |
| 5E-03 | 1.18 | 0.090 | 6.74E-02 | 0.032 | 1.18 | 0.098 | 9.32E-02 | 0.211 |
| 5E-02 | 1.25 | 0.091 | 1.23E-02 | 0.034 | 1.17 | 0.097 | 1.07E-01 | 0.211 |
| 1 | 1.30 | 0.091 | 3.33E-03 | 0.036 | 1.26 | 0.097 | 1.56E-02 | 0.214 |

**Average of the significant models (regression coefficient p-value < 0.05)**
| PRS | Average<br>OR | Average<br>R<sup>2</sup> | Null R<sup>2</sup> ** | Incremental R<sup>2</sup>***<br>over PRS null model |
| --- | --- | --- | --- | --- |
| EA | 1.13 | 0.044 | 0.004 | 0.041 |
| MA | 1.14 | 0.039 | 0.004 | 0.036 |
| AA | 1.25 | 0.034 | 0.004 | 0.030 |

| **PPRS** | Average OR | Average $R^2$ | PPRS Null $R^2$ ** | Incremental $R^{2***}$ over null model | Incremental $R^{2***}$ over PPRS null model |
| --- | --- | --- | --- | --- | --- |
| EA | 1.12 | 0.231 | 0.193 | 0.228 (61.5-fold) | 0.038 (19.6%) |
| MA | 1.13 | 0.228 | 0.201 | 0.224 (58.8-fold) | 0.027 (13.2%) |
| AA | 1.28 | 0.215 | 0.193 | 0.211 (57.0-fold) | 0.021 (11.0%) |

OR = odds ratio; SE = standard error; $R^2$ = psuedo- $R^2$
\* PRS: PRS + basic covariates [Model(1) = PCOS ~ PRS + PC1-4 + Site ]
\* PPRS: PRS + PCOS component phenotypes + basic covariates [PPRS = PCOS ~ PRS + PC1-4 + Site + Hirsutism + Female Infertility + Irregular Menses]
\*\* Null model: basic covariates only [Null Model = PCOS ~ PC1-4 + Site]
\*\* PPRS Null model: PCOS component phenotypes + basic covariates [PPRS Null Model = PCOS ~ PC1-4 + Site + Hirsutism + Female Infertility + Irregular Menses]
\*\*\* Improvement rate: (Incremental change in pseudo- $R^2$ between the model with PRS/PPRS and the null model without PRS/PPRS) / (Pseudo- $R^2$ of the null model without PRS/PPRS)

The regression models built in the training set were then used to predict PCOS case status in the testing dataset. A model including PRS yielded average prediction accuracy of 0.55, sensitivity of 0.55, specificity of 0.76 with an average area under the receiving operating characteristic curve (AUC) of 0.72 in the EA participants (Table 3).

**Table 3:** Average performance of PRS prediction algorithms in the female cohorts of European (n=33,869), ancestry (n= 49,365) and African (n=8,198) participants.

**Summary - Average**
| **PRS\*** | **Accuracy** | **Sensitivity** | **Specificity** | **Balanced Accuracy\*\*\*** | **AUC\*\*\*\*** |
| --- | --- | --- | --- | --- | --- |
| **European (n=33,869)** | 0.551 | 0.547 | 0.755 | 0.651 | 0.715 |
| **Multiancestry (n= 49,365)** | 0.533 | 0.529 | 0.736 | 0.632 | 0.693 |
| **African (n=8,198)** | 0.496 | 0.494 | 0.590 | 0.542 | 0.543 |
| **PPRS\*\*** | **Accuracy** | **Sensitivity** | **Specificity** | **Balanced Accuracy\*\*\*** | **AUC\*\*\*\*** |
| **European (n=33,869)** | 0.873 | 0.876 | 0.717 | 0.797 | 0.870 |
| **Multiancestry (n= 49,365)** | 0.881 | 0.886 | 0.640 | 0.763 | 0.823 |
| **African (n=8,198)** | 0.864 | 0.872 | 0.522 | 0.697 | 0.706 |

\* PRS: PRS + basic covariates [PRS Model = PCOS ~ PRS + PC14 + Site ]
\*\* PPRS: PRS + PCOS component phenotypes + basic covariates [PPRS Model = PCOS ~ PRS + PC1-4 + Site + Hirsutism + Female Infertility + Irregular Menses]
\*\*\* Balanced Accuracy = (Sensitivity + Specificity)/2
\*\*\*\* AUC = Area Under the Curve

#### B. Stratification ability of each prediction model with different PRS

The percentage of PCOS-diagnosed patients increases in higher PRS quantiles, and the individuals in the highest PRS group tend to have higher average BMI. In the genome-wide PRS calculation with SNVs with p-value ≤ 1, the average BMI of the individuals in highest PRS quantile is 1.1 kg/m^2^ higher than the individuals in the lowest PRS group (Cohen’s d=0.16, t-test p-value=1.06×10^-9^) (Figure 2 **and** Table 4). The finding confirms the positive correlation between elevated generic risk for PCOS, actual PCOS diagnosis, and higher risk for increased BMI.

**Fig 2.**
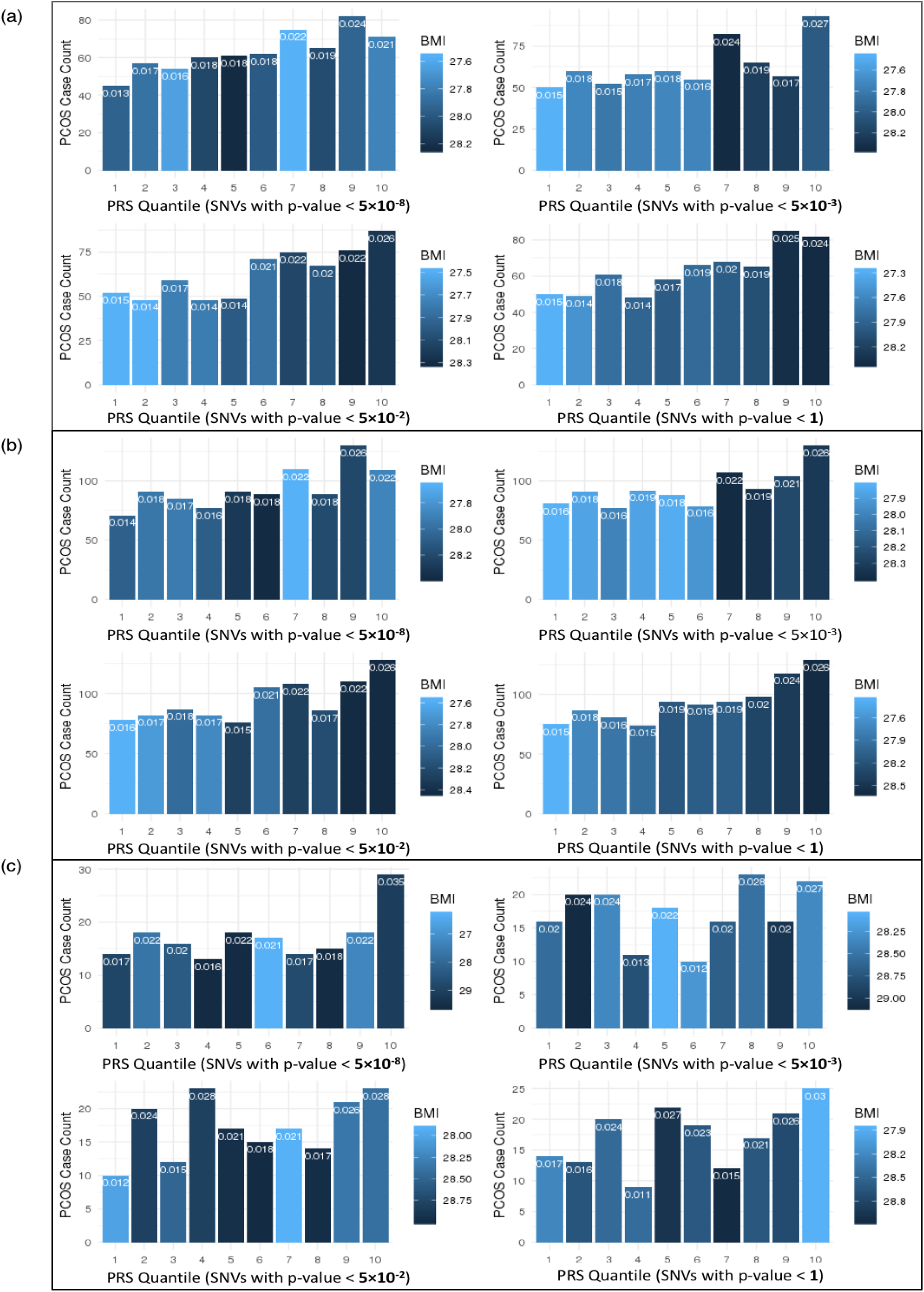
**Stratification performance by quantile of PRS models**, including PCs 1-4 and site as covariates, in (a) EA, (b) MA, and (c) AA populations. Group 1 includes those with the lowest PRS, and group 10 includes those with the highest. Bar colors indicate the average BMI in the quantile (darker indicates higher BMI), while the proportion of PCOS-diagnosed patients in each group is indicated at the top of each bar.

**Table 4:** Quantile analysis of PRS in the female European cohort (n=33,869) (PRS SNVs’ p-value<5×10^-8^ and p-value≤1 only).

|  | GROUP* | PCOS cases | PCOS prop** | Average BMI (kg/m<sup>2</sup>) | Average PRS |
| --- | --- | --- | --- | --- | --- |
| **PRS**<br>**P < 5×10<sup>-8</sup>** | 1 | 45 | 1.3% | 27.9 | -1.750 |
|  | 2 | 57 | 1.7% | 27.9 | -0.950 |
|  | 3 | 54 | 1.6% | 27.6 | -0.813 |
|  | 4 | 60 | 1.8% | 28.1 | -0.239 |
|  | 5 | 61 | 1.8% | 28.2 | -0.068 |
|  | 6 | 62 | 1.8% | 28.0 | 0.014 |
|  | 7 | 75 | 2.2% | 27.6 | 0.248 |
|  | 8 | 65 | 1.9% | 28.1 | 0.810 |
|  | 9 | 82 | 2.4% | 27.9 | 0.952 |
|  | 10 | 71 | 2.1% | 27.8 | 1.790 |
| ... |  |  |  |  |  |
| **PRS**<br>**P ≤ 1** | 1 | 50 | 1.5% | 27.3 | -1.790 |
|  | 2 | 49 | 1.5% | 27.5 | -1.020 |
|  | 3 | 61 | 1.8% | 27.8 | -0.654 |
|  | 4 | 48 | 1.4% | 27.9 | -0.369 |
|  | 5 | 58 | 1.7% | 28.0 | -0.113 |
|  | 6 | 66 | 2.0% | 27.8 | 0.132 |
|  | 7 | 68 | 2.0% | 28.0 | 0.386 |
|  | 8 | 65 | 1.9% | 28.1 | 0.672 |
|  | 9 | 85 | 2.5% | 28.4 | 1.040 |
|  | 10 | 82 | 2.4% | 28.4 | 1.720 |

PRS is adjusted with covariates and scaled for standardization.
\* Higher group number indicates higher PRS
\*\* Proportion of PCOS case patients in the quantile

The subsequent t-test reveals that PRS of case patients are significantly higher than the controls in all the nominally significant PRS models with regression p-value < 0.05, implying that higher genetic risk scores indicate higher occurrence of PCOS diagnosis (p-value=2.15×10^-4^, 7.75×10^-4^, 2.43×10^-4^, 2.51×10^-5^, 3.12×10^-5^ in PRS model SNVs’ p-value < 5×10^-8^, 5×10^-4^, 5×10^-3^, 5×10^-2^, 1 respectively) (**Supplementary Table 2**).

#### C. Performance improvement by the PRS variable

All the PRS models containing PCOS-PRS provided an improved fit over the null model by increasing the estimated explained sum of squares (pseudo-R^2^) by McFadden’s [37]. The average increase of pseudo-R^2^ by the PRS variable in EA samples is 0.040, which is a 10-fold improvement (=0.040/0.004) over the null model. The ANOVA p-values of differentiating the PRS models from the null model are all under 1×10^-31^, which validate the statistical significance of the performance improvement over the null model (Table 2 **and Supplementary Table 3**).

### Evaluation of PRS in multi-ancestry and African ancestry participants

#### A. Predictive ability of each prediction model with different PRS

In the training set of the MA cohort (n=49,365 with 949 PCOS cases), the coefficient p-values of all PRS variables remain significant with positive beta coefficients (Table 2**; model1**). The average OR of PRS is 1.14 (average SE=0.038) and the average pseudo-R^2^ value is 0.039, indicating that 3.9% of the phenotypic variance in the MA cohort could be explained by the PRS model. In the training set of AA participants (n=8,198 with 172 PCOS cases), the coefficient p-values of PRS variables remain overall significant except for two PRS models of SNVs with p-value < 5×10^-8^ and p-value < 5×10^-3^ which may be due to the smaller sample size. Even though the regression p-values of the PRS variable do not show uniform performance in AA as compared to EA, the nominally significant PRS models generate a higher effect size in the AA samples compared to the other ancestry groups. The average OR of PRS models in the AA is 1.25 (SE=0.089), higher than both the EA (OR=1.13) and MA (OR=1.14). This is possibly due to the low RAF of PCOS risk variants in AA compared to EA (**Supplementary Table 1**).

For the testing dataset, PRS prediction displays an average 0.533 of accuracy, 0.529 of sensitivity, 0.736 of specificity with an average AUC of 0.693 in the multi-ancestry cohort. The out-of-sample performance in AA yielded an average AUC of 0.543 and showed an overall lower average accuracy (0.496), sensitivity (0.494) and specificity (0.590) compared to other ancestry groups (Table 3).

#### B. Stratification ability of each prediction model with different PRS

In the MA cohort, the proportion of PCOS patients increases from 1.5% in the lowest quantile to 2.6% in the highest quantile in the PRS calculation of SNVs with p-value ≤ 1. The average BMI of the participants in the highest PRS quantile is 1.2 kg/m^2^ higher (Cohen’s d=0.17, t-test p-value=1.62×10^-13^) than the participants in the lowest PRS group (**Supplementary Table 4**, Figure 2(b)).

In the AA cohort, the number of PCOS patients does not always increase with higher PRS quantile, but the observation of an excess of PCOS patients in the highest PRS quantile is generally consistent across the models (Figure 2c). In the full-inclusive PRS model (SNVs with p-value ≤ 1), the rate of PCOS patients increases from 1.7% in the lowest quantile to 3.1% in the highest PRS quantile **(Supplementary table 4).** The observed increase of the rate of PCOS patients is most pronounced in the PRS model with genome-wide significant variants (SNVs with p-value < 5×10^-8^), as the PCOS case rate doubles from 1.7% in the lowest quantile to 3.5% in the highest PRS quantile. We did not identify any notable trends in BMI in AA participants, which is depicted by the quantile color changes in Figure 2(c).

An independent t-test confirms the significant differences of average PRS between PCOS cases and controls in MA across the models. The PRS difference between PCOS MA cases and controls is 0.165 after scaling with a full-inclusive PRS model, SNVs with p-value ≤ 1 (Cohen’s d=0.201, t-test p-value=2.62×10^-6^). In AA, only the full-inclusive PRS model shows statistically significant difference between PCOS cases and controls with a PRS difference of 0.175 (Cohen’s d=0.191, t-test p-value=2.90×10^-2^) **(Supplementary Table 2).**

#### C. Performance improvement by the PRS variable

In the joint ancestry participants, all the prediction models containing the PRS variable provide a better fit over the null model by increasing the average pseudo-R^2^ to 0.039, which is an 8.75-fold increase (=0.035/0.004) in explanatory power (Table 2). The subsequent ANOVA analysis confirms the statistical significance of the improved fits over the null model with all p-values<1×10^-46^ **(Supplementary table 3)**.

In the AA samples, the statistically significant PRS models show the average pseudo-R^2^ of 0.034, which has the poorest fit among the ancestries. The models show an average pseudo-R^2^ improvement of 7.5-fold increase (=0.030/0.004) from the null model without PRS (Table 2). Even with the lowest average incremental pseudo-R^2^(0.030) among the ancestries, the significant difference between the PRS models and the null model in Africans are confirmed with all ANOVA p-values<5×10^-3^ **(Supplementary table 3)**.

### Development of PPRS prediction algorithms with PRS and PCOS component phenotypes

The addition of PCOS component EHR phenotypes to polygenic risk prediction significantly improved the predictive accuracy (Table 2; **model2 and** Figure 3). The average pseudo-R^2^ of the PPRS is 0.231 in EA, 0.228 in MA, and 0.215 in AA samples, which indicates an average 14.7% improvement in pseudo-R^2^ (19.6% in EA, 13.2% in AA, 11.0% in MA) over the PPRS null model by the inclusion of PCOS component phenotypes. Compared to the basic null model, the PPRS prediction boosts the average predictive performance (pseudo-R^2^) by approximately 60 times (61.5-fold in EA, 58.8-fold in AA, 57.0-fold in MA) by the combinational use of PCOS component EHR phenotypes and PRS. Of note, the PRS variable’s p-values in every PPRS model remain consistently valid in the MA samples (p-values<5×10^-3^), whereas it was not always significant in AA or even EA samples. The ORs of the PRS and PPRS remain similar across the ancestries (Figure 4).

**Figure 3.**
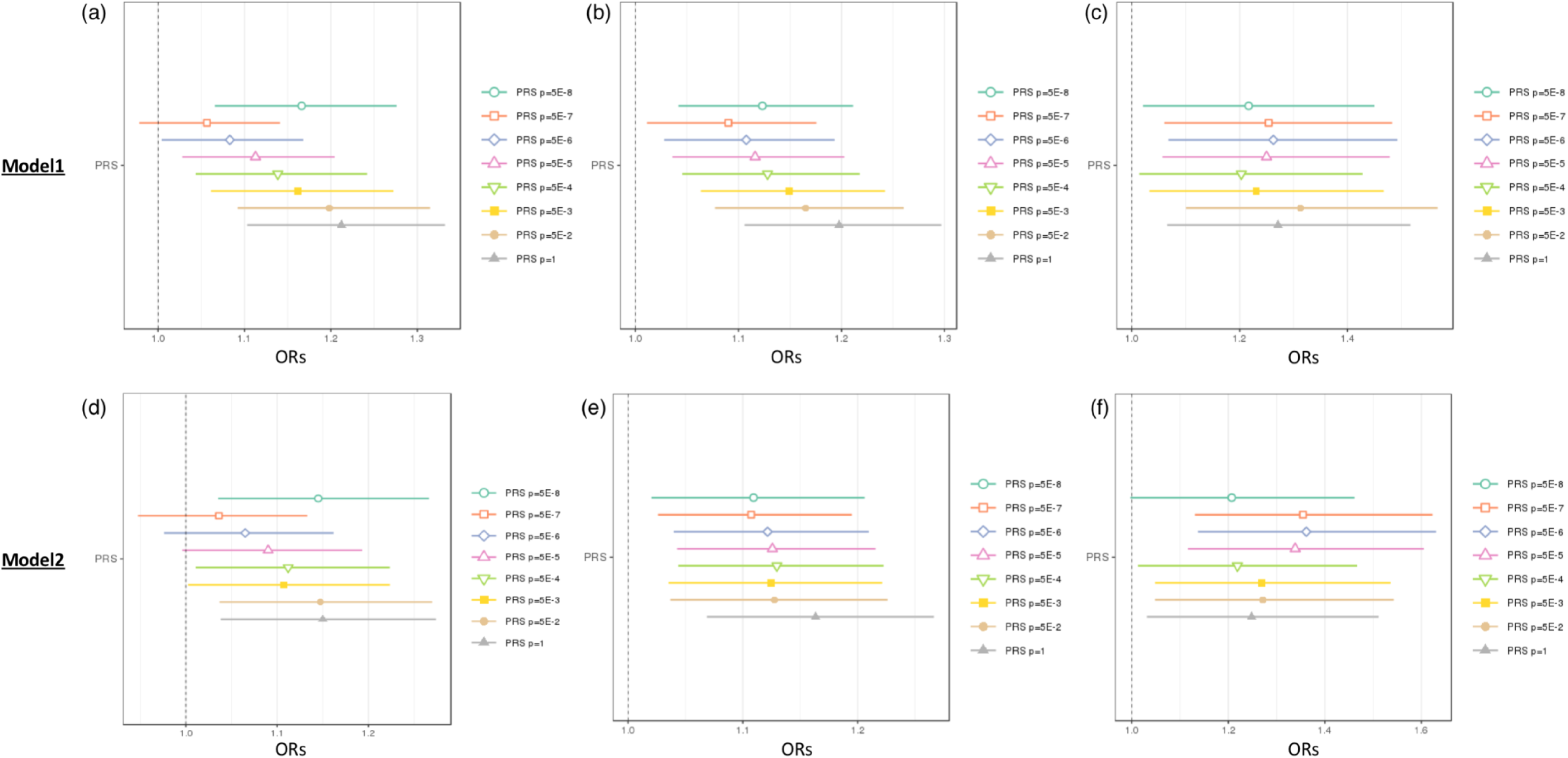
Comparison of odds ratios (ORs) for the PRS and PPRS in (a) EA, (b) MA, and (c) AA cohorts, at different PRS/PPRS inclusion thresholds by GWAS p-value. The top row shows OR distributions for the PRS model, which adjusted for basic covariates [PRS Model = PCOS ∼ PRS + PC1-4 + Site]. The bottom row shows OR distributions for the PPRS model which adjusted for the same basic covariates as well as PCOS EHR component phenotypes [PPRS Model = PCOS ∼ PRS + PC1-4 + Site + Hirsutism + Female Infertility + Irregular Menses].

**Figure 4.**
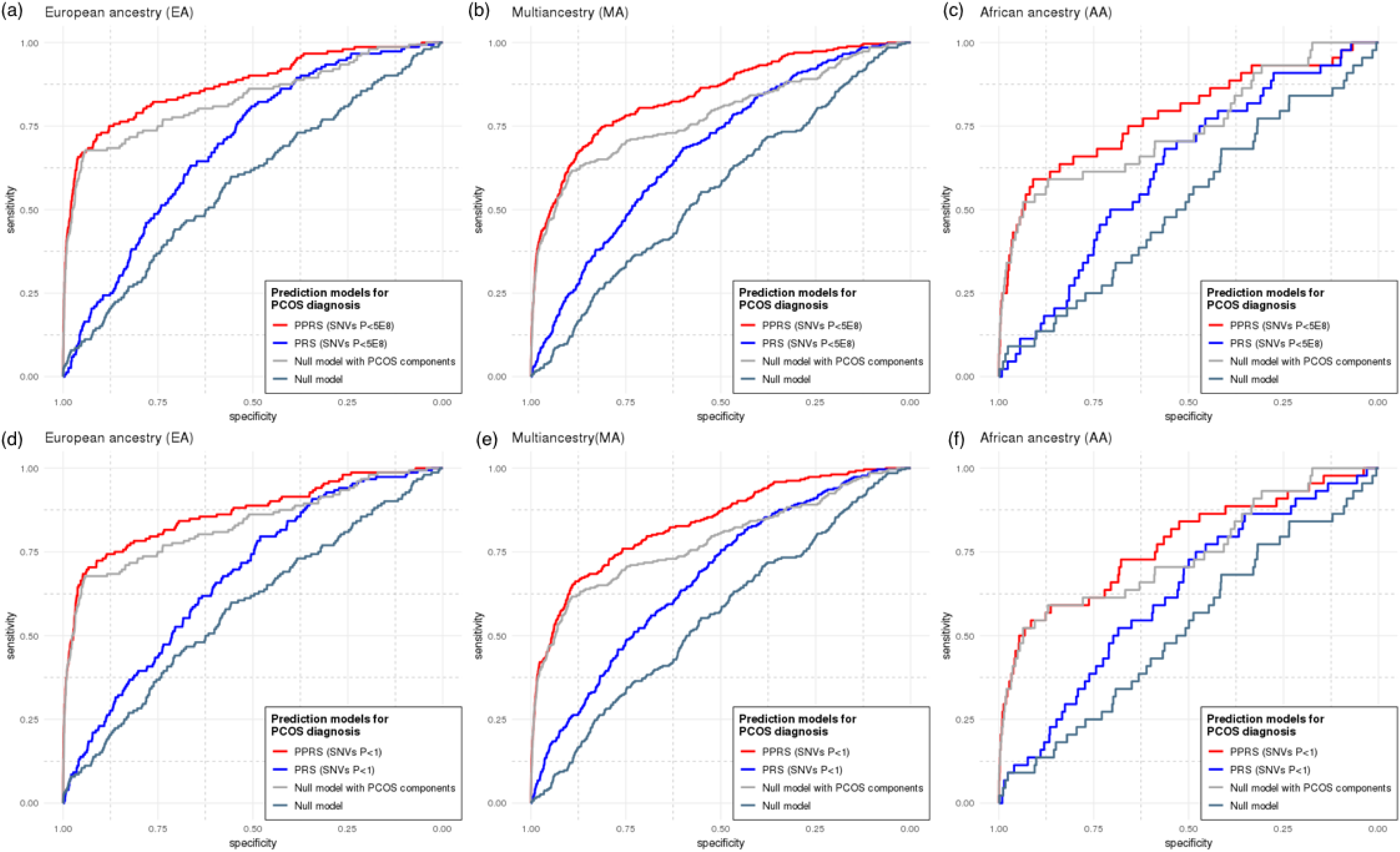
Comparison of Receiving Operating Curves (ROC) of the PPRS and PRS prediction models for PCOS diagnosis. The models with the genome-wide significant SNVs (p-value < 5×10^-8^) were evaluated in females of (a) EA, (b) MA, and (c) AA cohorts, along with the full-inclusive prediction models (p-value < 1) in females of (d) EA, (e) MA, and (f) AA cohorts. The areas under the curve (AUC) are provided in Table 2 and Supplementary Table 2. PRS model adjusted for basic covariates [PRS Model = PCOS ∼ PRS + PC1-4 + Site], and PPRS model adjusted for the same basic covariates as well as PCOS EHR component phenotypes [PPRS Model = PCOS ∼ PRS + PC1-4 + Site + Hirsutism + Female Infertility + Irregular Menses]. Null models only included the basic covariates without the PRS variable.

The subsequent ANOVA tested that all the pairs between PPRS and PPRS null models were statistically distinct across the cohorts and every PPRS model show the improved fit over the PPRS null model **(Supplementary Table 3)**. The average ORs of irregular menstruation (ICD9 code 626.4, ICD10 code N92.6), female infertility (ICD9 code 627, ICD10 code N97.0) and hirsutism (ICD9 code 704.1, ICD10 code L68.0) for PCOS prediction were, as expected, strong across the cohorts: 5.49, 10.9, and 17.1, respectively **(Supplementary Table 5)**.

### Clinical phenome analysis

#### A. Associated phenotypes *with PRS **(PheWAS-1)***

The general scheme of our PheWAS analyses are depicted in Figure 5a. Based on the model examination described above, the genome-wide PRS that includes all SNVs with p-value ≤ 1 was selected as the best performing PRS model and used for phenome-wide analysis. The phenomes of 49,343 female participants and 41,669 male participants were analyzed separately to test for association with high genetic risk for PCOS.

**Fig 5.**
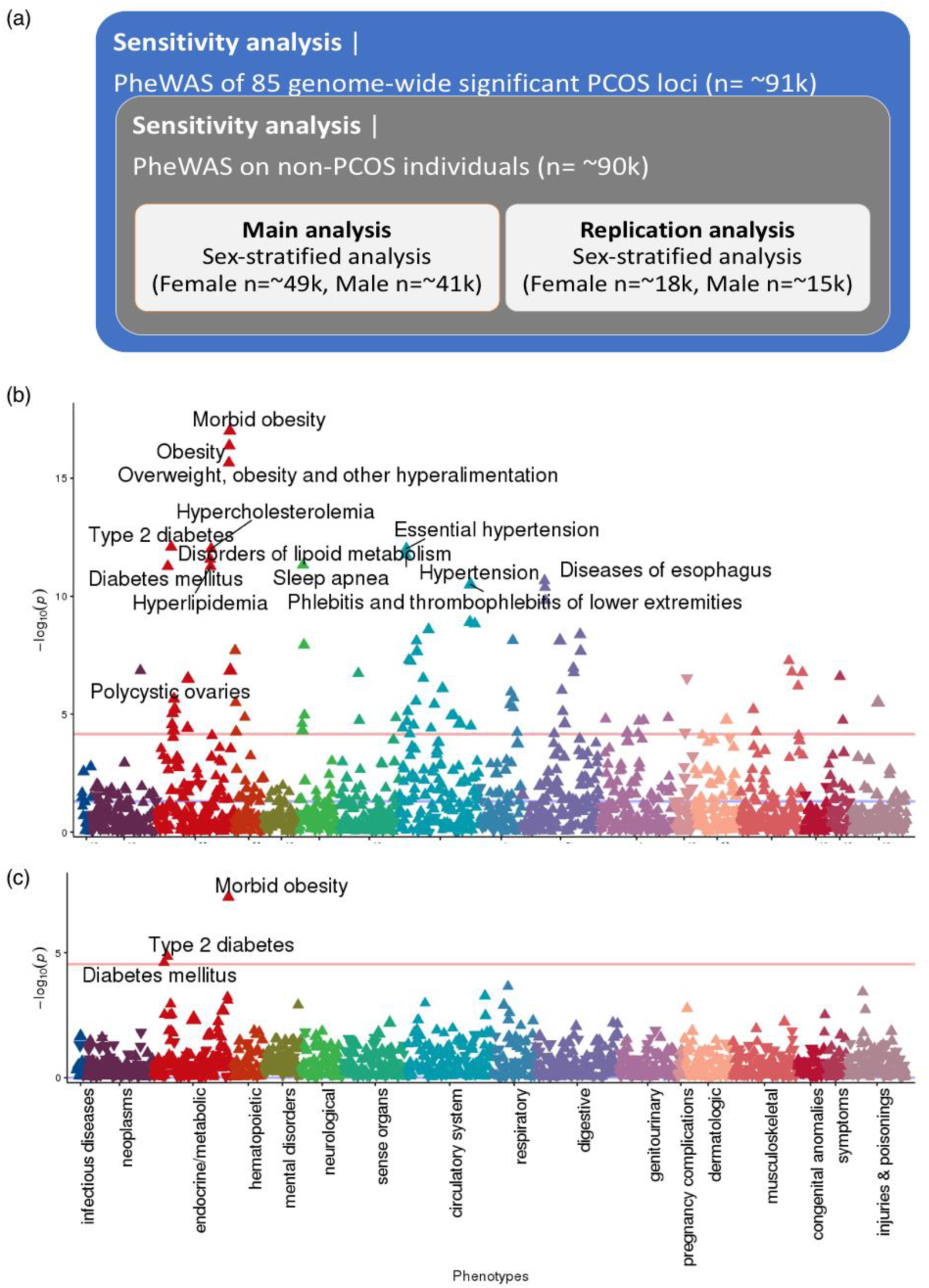
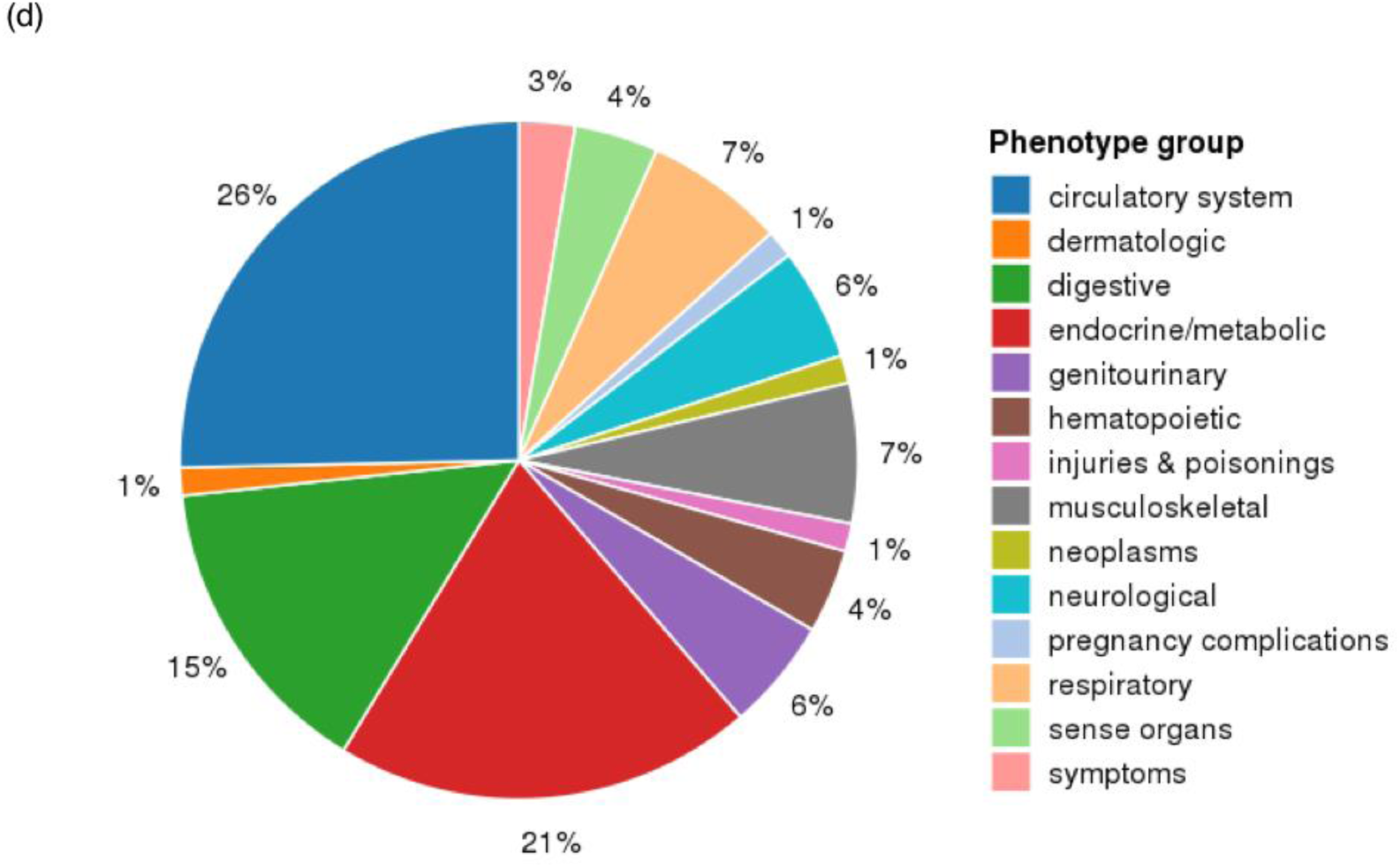
PheWAS scheme and results using PRS. (a) PheWAS scheme and sample sizes; (b) PheWAS Manhattan plot of PRS (SNVs with p-value ≤ 1); (c) PheWAS Manhattan plot of PRS (SNVs with p-value < 5E-08); (d) pie chart summarizing PheWAS groups. In Manhattan plots (b) and (c), the x-axis represents the EHR phenotype categorical group and the y-axis represents the negative log(10) of the PheWAS p-value. Red lines indicate the cutoff for phenome-wide significance. For readability, only the most significant associations are annotated. Full lists of phenome-wide significant results are provided in Supplementary Tables 5 and 6, respectively. The pie chart in (d) shows EHR categories for the 72 phenome-wide significant phenotypes identified through PheWAS of the genome-wide PRS (SNVs with p-value ≤ 1).

**Table 5:** (a) 16 significant phenotypes of PRS (SNVs’ p-value ≤ 1) female-stratified PheWAS that were phenome-wide significant in the discovery cohort (n=49,343) and ssfully replicated in the independent VU cohort (n=18,096). (b) 3 phenome-wide significant results of PCOS-PRS (SNPs with P < 1) male-stratified PheWAS from the ery cohort (n=41,669) and replication cohort (n=15,612)

| (a) |  |  | Discovery analysis |  |  |  |  | Replication analysis |  |  |  |  |
| --- | --- | --- | --- | --- | --- | --- | --- | --- | --- | --- | --- | --- |
| phecode | description | group | OR | SE | p | n_total | n_cases | OR | SE | p | n_total | n_cases |
| 278.11 | Morbid obesity | endocrine/metabolic | 1.010 | 0.001 | 9.74E-18 | 37108 | 6790 | 1.116 | 0.029 | 1.64E-04 | 15329 | 1762 |
| 278.1 | Obesity | endocrine/metabolic | 1.008 | 0.001 | 4.14E-17 | 44267 | 13949 | 1.087 | 0.022 | 1.29E-04 | 17051 | 3484 |
| 278 | Overweight, obesity and other hyperalimentation | endocrine/metabolic | 1.007 | 0.001 | 2.20E-16 | 47803 | 17485 | 1.077 | 0.020 | 1.44E-04 | 18096 | 4529 |
| 250.2 | Type 2 diabetes | endocrine/metabolic | 1.007 | 0.001 | 8.18E-13 | 42874 | 10800 | 1.081 | 0.022 | 3.70E-04 | 16562 | 3660 |
| 327.3 | Sleep apnea | neurological | 1.008 | 0.001 | 4.71E-12 | 40673 | 6503 | 1.096 | 0.028 | 1.33E-03 | 15602 | 1847 |
| 250 | Diabetes mellitus | endocrine/metabolic | 1.007 | 0.001 | 5.39E-12 | 43325 | 11251 | 1.079 | 0.021 | 3.56E-04 | 16763 | 3861 |
| 571 | Chronic liver disease and cirrhosis | digestive | 1.008 | 0.001 | 4.17E-09 | 40531 | 4582 | 1.093 | 0.032 | 4.64E-03 | 15369 | 1463 |
| 539 | Bariatric surgery | digestive | 1.012 | 0.002 | 7.59E-09 | 47803 | 2034 | 1.202 | 0.055 | 8.00E-04 | 18096 | 439 |
| 327.32 | Obstructive sleep apnea | neurological | 1.007 | 0.001 | 1.16E-08 | 39291 | 5121 | 1.098 | 0.032 | 3.98E-03 | 15138 | 1383 |
| 571.5 | Other chronic nonalcoholic liver disease | digestive | 1.008 | 0.001 | 2.13E-08 | 40251 | 4302 | 1.112 | 0.033 | 1.38E-03 | 15219 | 1313 |
| 743.9 | Osteopenia or other disorder of bone and cartilage | musculoskeletal | 1.005 | 0.001 | 1.71E-07 | 43335 | 11354 | 0.956 | 0.022 | 4.45E-02 | 16019 | 3263 |
| **256.4** | **Polycystic ovaries** | **endocrine/metabolic** | **1.015** | **0.003** | **3.16E-07** | **40696** | **942** | **1.174** | **0.069** | **1.93E-02** | **15637** | **281** |
| 743 | Osteoporosis, osteopenia and pathological fracture | musculoskeletal | 1.004 | 0.001 | 6.38E-07 | 47803 | 15822 | 0.957 | 0.019 | 1.84E-02 | 18096 | 5340 |
| 250.3 | Insulin pump user | endocrine/metabolic | 1.008 | 0.002 | 2.25E-06 | 35057 | 2983 | 1.136 | 0.036 | 3.42E-04 | 14065 | 1163 |
| 250.23 | Type 2 diabetes with ophthalmic manifestations | endocrine/metabolic | 1.010 | 0.002 | 9.20E-06 | 33663 | 1589 | 1.221 | 0.062 | 1.20E-03 | 13272 | 370 |
| 627 | Menopausal and postmenopausal disorders | genitourinary | 1.004 | 0.001 | 1.40E-05 | 40468 | 14392 | 0.947 | 0.020 | 7.64E-03 | 16061 | 4301 |

(b)
|  |  |  | <u>Discovery analysis</u> |  |  |  |  | <u>Replication analysis</u> |  |  |  |  |
| --- | --- | --- | --- | --- | --- | --- | --- | --- | --- | --- | --- | --- |
| **phecode** | **description** | **group** | **OR** | **SE** | **p** | **n_total** | **n_cases** | **OR** | **SE** | **p** | **n_total** | **n_cases** |
| 278.11 | Morbid obesity | endocrine/metabolic | 1.009 | 0.002 | 5.93E-08 | 32456 | 3489 | 1.049 | 0.036 | 1.78E-01 | 13465 | 1082 |
| 250.2 | Type 2 diabetes | endocrine/metabolic | 1.005 | 0.001 | 1.41E-05 | 36835 | 10984 | 1.031 | 0.021 | 1.49E-01 | 14000 | 4198 |
| 250 | Diabetes mellitus | endocrine/metabolic | 1.005 | 0.001 | 2.47E-05 | 37199 | 11348 | 1.029 | 0.021 | 1.70E-01 | 14180 | 4378 |

Phenome-wide significant threshold: p-value < 2.9E-5

In the female PheWAS with PRS, 75 EHR phenotypes were identified with phenome wide significance (Figure 5b, **Supplementary Table 6a)**. ‘Morbid obesity’ (phecode 278.11) and obesity-related endocrine phenotypes, including ‘overweight, obesity, and other hyperalimentation’ (phecode 278), ‘type 2 diabetes’ (phecode 250.2), ‘essential hypertension’ (phecode 401.1) ‘hypercholesterolemia’ (phecode 272.11), ‘hypertension’ (phecode 401), ‘disorders of lipid metabolism’ (phecode 272) are the top-ranked. The phenome-wide significant association of ‘polycystic ovaries’ (phecode 256.4) and PCOS-PRS are observed with one of the largest effect sizes (OR=1.015) among the result.

As a complex endocrine disorder, the PCOS pathophysiology seems to be tightly linked to the expression of endocrine or circulatory system manifestation. Among the 75 phenome-wide significant traits with PRS, the phenotypes of circulatory system (26.0%) and endocrine/metabolic system (21.0%) appeared the most frequently (Figure 5d), while the four highest associated phenotypes are all endocrine/metabolic features.

Among the remainder of the phenome-wide significant phenotypes, associations of musculoskeletal phenotypes like ‘osteoarthrosis’ (phecode 740 and 740.9) or ‘calcaneal spur; Exostosis NOS’ (phecode 726.4) possibly imply the hormonal changes on the skeletal system impacted by PCOS epidemiology. Multiple symptomatic genitourinary phenotypes of PCOS were also identified: ‘abnormal mammogram’ (phecode 611.1) or ‘other signs and symptoms in breast’ (phecode 613.7). An obesity-related pulmonary disorder of ‘sleep apnea’ (phecode 327.3) is also observed (classified as neurological phenotype in phecode map) with ‘obstructive sleep apnea’ (phecode 327.32). We could not identify any psychological or depression related phenotype that is known to have genetic correlation with the hormonal changes of PCOS.

The overall low range of OR (1.004∼1.010) of the PheWAS results should be noted, which is assumedly due to the aggregated effects of the low impact SNVs for PCOS, especially in the full-inclusive PRS with the entire GWAS SNVs. The ORs from the generic PheWAS of individual PCOS SNVs are observed to be higher before merging them into the cumulative PRS, which is described later **(Supplementary Table 7)**.

In the replication analysis on an independent cohort of 18,096 EA females (BioVU), 16 out of 75 phenome-wide signals from the discovery analysis are replicated including ‘PCOS’ (p-value=1.93×10^-2^, phecode 256.4) with the positive OR of 1.174 (Table 5a). Half of the replicated phenotypes (8 out of 16) belong to the endocrine/metabolic category. In particular, the following obesity-related endocrine phenotypes exhibit strong evidence of replication after multiple testing correction (p-value < 6.7×10^-5^, 0.05/75): ‘morbid obesity’ (phecode 278.11), ‘obesity’ (phecode 278.1), ‘overweight, obesity and other hyperalimentation’ (phecode 278). The well-known comorbidity between ‘type 2 diabetes’ (phecode 250.2) and PCOS is also identified along with other diabetic syndromes like ‘diabetes mellitus’ (phecode 250). Other notable replicated phenotypes included multiple neurological and digestive manifestations, which have well-known association to obesity, such as ‘chronic liver disease and cirrhosis’ (phecode 571), ‘bariatric surgery’ (phecode 539) and ‘other chronic nonalcoholic liver disease’ (phecode 571.5). An obesity-related pulmonary disorder of ‘sleep apnea’ (phecode 327.3) is also observed (classified as neurological phenotype in phecode map) with ‘obstructive sleep apnea’ (phecode 327.32).

In male-specific PheWAS with PRS (SNVs with p-value ≤ 1) model, three metabolic phenotypes reached phenome-wide significance in the discovery analysis: ‘morbid obesity’ (phecode 278.11), ‘type 2 diabetes’ (phecode 250.2), ‘diabetes mellitus’ (phecode 250) which are known risk factors and/or co-morbidities for PCOS (Figure 5b, Table 5b, **Supplementary Table 6b)**. However, none of the associations were replicated in the replication analysis on 15,611 independent males. It is possible that the replication sample remained underpowered and larger sample sizes will be needed to distinguish these results from a true null result.

#### B. Sensitivity analysis – Case-excluded analysis **(PheWAS-2)**

After removing 949 PCOS patients in PheWAS investigation, we still identified 68 PRS-phenotype associations that reached phenome-wide significance **(Supplementary table 8)**, which is not very different from PheWAS-1. The result might be due to the challenge of current diagnosis practices in identifying PCOS cases, which implies the control groups are not completely excluding PCOS patients and possibly include some mixed signals from the unidentified PCOS cases. Alternatively, it is possible that genetic risk for PCOS remains a robust risk factor for these phenotypes even in the absence of clinical manifestations of PCOS.

The representative signals of diabetes/obesity-related endocrine traits that are identified in PheWAS-1 remained significant: ‘morbid obesity’ (phecode 278.11), ‘type 2 diabetes’ (phecode 250.2), ‘obesity’ (phecode 278.1), ‘overweight, obesity and other hyperalimentation’ (phecode 278), ‘diabetes mellitus’ (phecode 250), ‘hypercholesterolemia’ (phecode 272.11), ‘disorders of lipid metabolism’ (phecode 272) and ‘hyperlipidemia’ (phecode 272.1) etc.

Four phenotypes no longer remained phenome-wide significant in PheWAS-2 compared to PheWAS-1, including ‘menopausal and postmenopausal disorders’ (phecode 627), ‘iron deficiency anemias, unspecified or not due to blood loss’ (phecode 280.1), ‘sleep disorders’ (phecode 327) and ‘Insomnia’ (phecode 327.4). A new metabolic phenotype of ‘disorders of fluid, electrolyte, and acid-base balance’ (phecode 276) was phenome-wide significance in PheWAS-2 compared to PheWAS-1, but the association did not remain significant in replication analysis. The phenome-wide significant phenotype with the largest effect size in PheWAS-2 is ‘localized adiposity’ (OR=1.014, phecode 278.3), same as for PheWAS-1. It should be of note that the range of OR is low in PRS-PheWAS due to the cumulative effect sum of all PCOS susceptibility loci including low-effect variants.

#### C. Sensitivity analysis – Associations with individual PCOS susceptibility loci **(PheWAS-3)**

In the individual PheWAS of 85 PCOS genome-wide significant variants, even though no association survives phenome-wide significance, likely due to the multiple testing burden, 11 PCOS variants show notable association to ‘polycystic ovaries’ across the ancestry groups (Most significant variant hg19 chr11:30226528, OR=1.36, phecode 256.4), ranked as the second most significant phenotype **(Supplementary table 7)**. Out of top 100 associations in PheWAS-3, the largest number of associations were related to circulatory system for ‘thrombotic microangiopathy’ (31.0%). Endocrine/metabolic related phenotypes were the second most frequent category (21.0%) composed of either ‘PCOS’ or ‘ovarian dysfunction’, and 12% of the top associations were digestive traits, largely devoted to diverticular diseases. We did not identify any associations related to obesity or diabetes, which were the most significant phenotypic features found in PheWAS-1 and PheWAS-2.

## Discussion

A key question in precision medicine is how to identify patients at high risk for a given disease for the goal of targeting preventive care. In this study, we examined the ability of PRS to predict PCOS clinical diagnosis and mine comorbid EHR phenotypes with the ultimate goal of improving diagnostic accuracy for PCOS. We show that a PRS for PCOS can be used (a) to identify patients at elevated risk of PCOS and (b) to determine the comorbid or pleiotropic phenome-wide expression associated with PCOS in a clinical setting.

The primary accomplishment of this study is a systematic enhancement of the polygenic risk prediction by integration of additional disease component phenotypes in the EHR into a PPRS. The onset of hirsutism, menstrual dysfunction, or female infertility are representative symptoms of PCOS and essential in determining clinical hyperandrogenism [10, 40, 41]. They are not required for a diagnosis of PCOS per se, but are useful in suggesting PCOS in a clinical context. The PPRS significantly improves the average explanatory power (pseudo-R^2^) of PCOS prediction by 0.221 (59.1-fold increase) compared to the null model without PRS or component phenotypes, and by 0.037 (14.7% increase) over the null model with the component phenotypes alone (Table 2 **and** Figure 4). In contrast to the previous studies that attempted to identify PCOS diagnosis with risk score calculation [13, 42], our algorithm did not limit risk predictor in a single-dimension, using both phenotype and genotype markers with polygenic inheritance, and extensively demonstrated the predictive performance of PPRS with several machine-learning techniques. The findings shown here strengthen the potential clinical utility of PPRS as a disease predictor, particularly when combined with component symptom information available within the EHR.

To date, research has consistently shown that the PRS built from EA GWAS data does not perform as robustly across non-EA samples. In this study, we assessed the performance of a Eurocentrically built PCOS-PRS on the samples of EA, AA, and the joint MA cohorts. Undeniably, validation statistics varied by ancestry group and the PCOS diagnosis prediction in AA cohort shows the poorest performance. However, it is of note that more than half of the tested models in AA still show statistical significance in terms of regression p-value, and those models display a reliable efficiency for PCOS detection in effect size and AUC (Table 3). Interestingly, the ORs for PRS differ across the ancestry cohorts, and somewhat higher in some prediction models in AA (average OR of model1=1.25, model2=1.28) and MA samples (average OR of model1=1.14, model2=1.13) than EA samples (average OR of model1=1.13, model2=1.12). The overall ORs of the PRS variable are fairly stable throughout all polygenic prediction models (OR 1.12∼1.28). The observed significance of the PRS variable in the MA cohort, more stable than in the EA or AA participants alone, is likely due to the increased statistical power with larger sample size that counters the sample heterogeneity introduced. In addition, we found that the accumulation of genetic variants did not always increase the predictive capability of PRS in terms of pseudo-R^2^ and OR (Figure 3, Table 2). This might be due to the different RAF of PCOS risk variants by different PRS p-value cutoffs, and the varying LD structure of the ancestry groups. Previous research has confirmed that the LD pattern varies between EA and Chinese women at the PCOS susceptibility loci encoding LH/choriogonadotropin receptor (*LHCGR*) and FSH receptor (*FSHR*) genes, but the reproducible signals of the loci are consistently associated to PCOS regardless of ancestry[43, 44]. Our sensitivity analysis (PheWAS-3) also suggests the varying phenotypic effect of PCOS loci in different ancestries, but confirms the strong association with PCOS nonetheless. These findings demonstrate the primary role of PCOS-PRS in cumulatively explaining substantial variation of disease susceptibility across ancestries even with differing LD structures, and extend the general utility of PPRS in disease prediction.

Furthermore, our PRS-based phenome-wide analysis revealed several clinical associations that are tightly linked with obesity, confirming the shared metabolic pathways between PCOS and obesity in a phenomic aspect. As obesity is a common finding which can be found in 50-65% of PCOS patients[10], and previous Mendelian randomization study revealed the causal relationship of BMI on PCOS etiology[45], many of our findings could be interpreted as phenotypic evidence of co-morbid obesity. ‘Morbid obesity’ (phecode 278.11), ‘hypercholesterolemia’ (phecode 272.11), ‘disorders of lipoid metabolism’ (phecode 272), ‘hyperlipidemia’ (phecode 272.1), ‘hypertension’ (phecode 401) or ‘abnormal glucose’ (phecode 250.4) are easily understandable with the context of heightened metabolic risks for obesity. ‘Sleep apnea’ (phecode 327.3) and ‘chronic liver disease and cirrhosis’ (phecode 571), ‘GERD’ (phecode 530.11), ‘diseases of esophagus’ (phecode 530 and 530.1) are either neurological, pulmonary or digestive assorted symptoms that are commonly found in the patients with obesity.

It is also noteworthy that there were 75 significant associations identified in women while in men, there were only three significantly associated diagnosis (morbid obesity, type 2 diabetes, diabetes mellitus) despite a similar sample size for males and females in the analysis. It is possible that the clinical consequences of high androgens in males are less likely to cause symptoms for which medical treatment is sought, or that these genetic variants only elevate androgen levels in a female ‘environment’ but not a male one. The three identified phenotypes in males additionally suggest that if an individual harbors high genetic risk for PCOS, the metabolic manifestations are similar regardless of sex.

Consistent with previous studies [13, 45], we identified phenotypic evidence of positive BMI association with genetic risk of PCOS. In the stratification analysis of PRS, our observation of the increased BMI in individuals with high risk of PCOS are evident in both EA and MA cohorts (Figure 2). The comorbid phenotypes could be driven by pleiotropy in which PCOS-associated genes also increase BMI, or could be due to under diagnosis of PCOS itself, in which case the association with obesity phenotypes may be a result of comorbidity with undiagnosed PCOS.

Several limitations to this study need to be acknowledged. First, the sample size of AA participants was relatively small which increases the likelihood of both false negative and false positive findings. Further investigation is needed to fully understand the overlap in PCOS genetic factors across multi-ancestry participants and the methodological application of Eurocentric PCOS-PRS to other genetic ancestries considering LD structure. Secondly, the phenotypic components we used for polygenic prediction are currently limited to only three representative phenotypes: hirsutism, irregular menstruation, and female infertility. Fueled by our PheWAS finding, the work could be extended by incorporating the additional phenotypes that might increase the likelihood of an eventual diagnosis. Also, the phecode of PCOS used for PheWAS was converted from ICD-9-CM 256.4 and ICD-10-CM E28.2, which was used as a proxy for capturing PCOS in the EMR. This phecode may not perfectly capture PCOS as they may or may not capture hyperandrogenemia. The selection bias in our discovery cohort should be acknowledged as well. Two of our participating sites (Geisinger and Marshfield) mainly recruited their patients for the study of obesity and type 2 diabetes, which resulted in a higher proportion of obese patients into their biobank and therefore may inflate the prevalence of PCOS in these subgroups. Lastly, due to the low diagnosis rate of PCOS patients in current EHR system, it is possible that unidentified PCOS cases could reduce power in each analysis.

Our approach has provided a novel methodological opportunity to stratify patients’ genetic risk and to discover the phenomic network associated with PCOS pathogenesis. Integrative analysis of the PRS-PheWAS enables the systematic interrogation of PCOS comorbidity patterns across the phenome, which cannot be readily identified by a single-variant approach. The identified phenomic networks could be used at the stage of first screening, prior to the testing of hormones or imaging of ovaries, or to help the patient and physician decide whether more extensive testing would be useful for PCOS diagnosis. From a precision medicine perspective, such an approach may provide a greater understanding of a patient’s clinical presentation and suspected diagnosis based on phenotypic or genetic variations.

## Acknowledgements

The phase III of the eMERGE Network was initiated and funded by the NHGRI through the following grants: U01HG008657 (Kaiser Permanente Washington/University of Washington School of Medicine); U01HG008685 (Brigham and Women’s Hospital); U01HG008672 (Vanderbilt University Medical Center); U01HG008666 (Cincinnati Children’s Hospital Medical Center); U01HG006379 (Mayo Clinic); U01HG008679 (Geisinger Clinic); U01HG008680 (Columbia University Health Sciences); U01HG008684 (Children’s Hospital of Philadelphia); U01HG008673 (Northwestern University); U01HG008701 (Vanderbilt University Medical Center serving as the Coordinating Center); U01HG008676 (Partners Healthcare/Broad Institute); and U01HG008664 (Baylor College of Medicine).

## Authors’ contributions

YYJ, LD and MGH designed the study; IBS and DRC imputed and quality controlled the genotype array data missing variants with input from GPJ; JAP, AOB, RC, DRC, JCD, DRVE, HH, JBH, SJH, KH, GPJ, FDM, SP, MDR, IBS contributed to eMERGE genotype and phenotype data generation; LD, MGH, FD, MJ, TK, CM generated PCOS GWAS data through the International PCOS consortium; YYJ performed statistical analysis in discovery cohort and validated the algorithms; KA performed statistical analysis in replication cohort; YYJ, ANK, LD and MGH interpreted the results; YYJ, KA, JAP, ANK, LD, MGH drafted the manuscript; YYJ designed the figures and created the tables; All authors critically reviewed the manuscript for important intellectual content; DRC, GPJ, MS and RLC obtained the funding.

